# Selective Biogenesis and Structural Diversity Shape tRNA-Derived Fragment Landscapes across Mouse Organs

**DOI:** 10.1101/2025.06.18.660468

**Authors:** Daisuke Ando, Sherif Rashad, Kuniyasu Niizuma

## Abstract

Transfer RNA derived small RNAs (tDRs) are increasingly recognized as versatile regulators, yet their physiological landscape remains poorly charted. We re-analyzed a small-RNA-seq dataset generated with tDR-optimized library preparation from seven adult mouse tissues to explore tissue specific tDRs enrichment. We catalogued 26901 unique nuclear tDRs (ntDRs) and 5114 mitochondrial tDRs (mtDRs). Library sizes and subclass composition varied markedly: heart was mtDRs-rich, whereas spleen produced the fewest reads but the greatest diversity. Clustering analysis segregated tissues, with spleen and lung forming a distinct immune cluster. Tissue-versus-all differential analysis showed the spleen harboring unique ntDRs and mtDRs. The brain preferentially accumulated 5′tiRNAs, while other organs were enriched for i-tRFs or tRF3s. Enriched fragments in different tissues originated from specific isoacceptors and isodecoder tRNAs independent of mature tRNA abundance, implying selective biogenesis rather than bulk turnover. G-quadruplex prediction revealed pronounced enrichment of potentially quadruplex-forming ntDRs in the spleen, predominantly Gly-, Asp- and Val-derived i-tRFs/tRF3s, suggesting structure-dependent functions in immune regulation. Collectively, our atlas reveals extensive tissue-specific heterogeneity in tDR biogenesis, sequence and structure, providing a framework for deciphering context-dependent regulatory roles of tDRs.

## Introduction

tRNA derived small non-coding RNAs (tDRs) are at the center of a growing research field that has grown exponentially in recent years^1,2^. Initially discovered as stress-induced Angiogenin-mediated cleavage products of tRNAs, yielding 5 prime and 3 prime tRNA halves (or 5’tiRNAs and 3’tiRNAs)^3–5^, tDRs have grown in complexity with many subclasses identified resulting from various modes of tRNA processing^6,7^. In addition to Angiogenin, several other RNAses have been reported to generate tDRs^6,8^. These enzymes act on specific tRNAs in specific contexts^9–12^. tRNA modifications also play important roles in regulating tDRs generation and function^13–16^. Thus, the production of tDRs is currently understood to be a highly ordered and regulated process and not a simple degradation product of tRNAs during cell death^1,5^. Nonetheless, our understanding of tDRs biogenesis remains incomplete, especially in physiological conditions.

Given the strong links between tDRs are cellular stress responses as well as many pathological conditions such as cancer, neurodegeneration, and metabolic disorders, most research on tDRs is done in pathologic cell states^2,17–22^. Thus, how tDRs shape gene expression and the proteome in physiological states remains understudied. Several studies have shown how tDRs regulate stem cell function and differentiation capacity^23,24^ and β-cell maturation^25^. In terms of how tDRs could regulate gene expression and the proteome, tDRs were shown to interact with RNA binding proteins (RBPs) to displace them from target mRNAs^26,27^. At the level of translation, tDRs were also shown to stimulate the translation of ribosomal mRNAs^28,29^. However, stress-induced 5’tiRNAs were shown to suppress translation initiation^3^. At the transcriptional level, tDRs were shown to act as miRNA-like molecules and to induce gene silencing^30^, although there is some debate regarding this view as tDRs can form secondary structures and G4 quadruplexes, thus do not fit the simplistic miRNA-like assumption^1,17,31^. 5’tiRNAs were shown to promote corresponding tRNA gene transcription^32^ while tRF3s (18∼22nt fragments originating from the 3’ side of the tRNA) were shown to regulate LTR-retrotransposons^33^. Thus, it appears that tDRs act via a variety of ways and are not restricted to one specific mode of activity. This heterogeneity in their function adds a layer of complexity to tDRs. tDR modifications, sequences, origins, site of production, secondary structure, and other factors could be contributing to the different ways by which a given tDR, or a set of tDRs, act^1,26,31,34,35^. Given the clear links between tDRs and diseases, either as potential therapeutic targets for cancers and other diseases or as biomarkers for various conditions^36–38^, it becomes clear that understanding their biogenesis and function is of utmost importance. However, no one approach is perfect, and thus combinations of biochemical, molecular, and computational approaches are necessary. Importantly, more high-quality datasets cataloging and exploring tDRs expression in various conditions and diseases are needed. Unfortunately, the current dependence on less optimal datasets, such as tDR sequences derived from The Cancer Genome Atlas (TCGA)^39^, in which the sequencing technique wasn’t optimized to capture neither tRNA nor tDRs especially in the context in the current accepted practices^40,41^, have caused some conflicting conclusions in terms of tDRs biogenesis and function. Thus, newer datasets with better techniques and coverage of tDRs are needed. One of the important approaches in studying non-coding RNAs is cataloging their expression in different cell states, tissues, and cell types. While this has been done effectively for microRNAs, for example, which are easier to sequence and have more robust over the shelf tools to prepare the libraries^42–44^, not much has been done in the tDR field using the same approach. While several databases have catalogued tDRs from hundreds of published sequencing datasets^45,46^, these databases did not include only datasets optimized for tDR detection, thus biases in tDR expression should be expected. Only one study systematically analyzed tDRs expression in different mouse tissues^42^. However, the library preparation for that study also did not include important steps as demethylation, deacetylation, and end-repair that are known to increase the coverage of tDRs and avoid biases from modifications and aminoacylation^40,47^. Thus, in this study we analyzed our previously published tRNA/tDR optimized small RNA sequencing dataset of 7 mouse tissues^48^. We aim to provide an in-depth understanding of the heterogeneity of tDRs across different tissues that could be linked to specific gene regulatory processes and tissue/cell physiology.

## Results

### Global patterns of tDRs expression in 7 mouse tissues

To explore the tissue specific expression of tDRs we used a previously published small RNA-seq dataset of seven different mouse tissues^48^. The reason for selecting this dataset and not others^42^ was the library preparation strategy. In the selected study^48^, small RNA fractions (<200nts) were deacylated, demethylated, and end repaired, which allow for a higher depth of coverage of tDRs^40,47^ as well as detection of mature tRNAs^48,49^. However, without such library preparation strategies, tDRs are usually underrepresented in libraries and their detection will be biased due to many factors such as RNA modifications or aminoacylation^40,47,49,50^.

To detect tDRs, we used tDRnamer^8^, which not only gives us the naming convention of the detected tDRs^51^, but also other information that allows for in-depth analysis. We analyzed the data using tDRnamer to detect nuclear encoded tDRs (ntDRs; those arising from nuclear encoded tRNAs) and mitochondrial tDRs (mtDRs) in separate runs. We filtered the detected tDRs to select between 15 and 50 nucleotides in length and have at least 10 counts in any sample for downstream analysis and created a custom classifier to classify the detected tDRs based on their start and end positions on mature tRNAs [Figure 1A-B]. Previously, it was shown that the absolute amounts of tRNAs differ between tissues, with spleen and lung being more enriched in other non-coding RNAs, reducing the reads mapped to tRNAs^48^. Analysis of the detected tDR reads revealed similar patterns to what was reported previously [Figure 1C-D], with the spleen having the lowest levels of detected ntDRs and mtDRs. The heart had the highest levels of detected mtDRs (more than three-fold the next tissue), also following what was reported previously^48^. Overall, we detected 26901 unique ntDRs and 5114 unique mtDRs.

**Figure 1:**
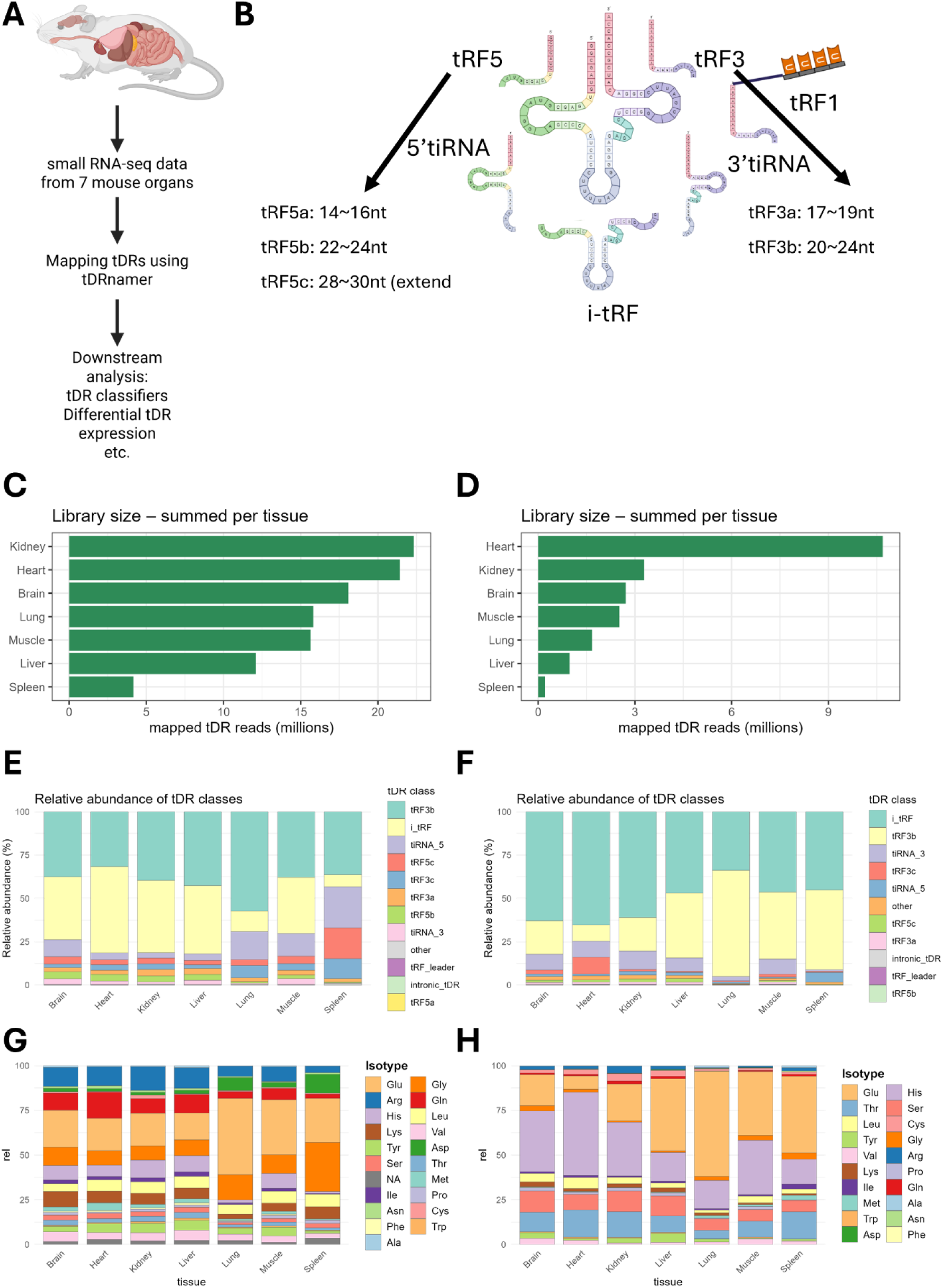
Global patterns of tDRs expression in mouse tissues. **A:** Strategy for analyzing tDRs in mouse tissues using tDRnamer. tRNA/tDR optimized small RNA sequencing dataset encompassing 7 mouse tissues (Brain, heart, lungs, kidneys, liver, spleen, and muscle, N = 3 per tissue)^48^ was used for the analysis. **B:** The known subtypes of tDRs used for the classifier in this study. **C:** Library size (in million reads) of the detected nuclear tDRs (ntDRs) per tissue (total number of mapped tDR reads that passed the filters and summed across samples). **D:** Library size of the detected mitochondrial tDRs (mtDRs) in different tissues. **E-F:** Relative distribution of different ntDRs (**E**) and mtDRs (**F**) subtypes/classes in different tissues. **G-H:** Relative distribution of isotype of origin (i.e. amino acids decoding tRNA) of ntDRs (**G**) and mtDRs (**H**) across tissues.

Examining the subclasses of tDRs, the most abundant in ntDRs were tRF3b, i-tRFs, and 5’tiRNAs (5’ halves or tiRNA_5 in the plot) [Figure 1E]. Indeed, variations between tissues were observed. For example, the spleen had the highest relative levels of 5’tiRNAs and tRF5c while the lungs had the lowest relative levels of i-tRFs and the highest relative levels of tRF3b fragments. In mtDRs, the dominant subclasses were i-tRFs and tRF3b, with variable levels between tissues but overall, more i-tRFs were detected in mtDRs compared to ntDRs [Figure 1F]. The isotype source of tDRs, i.e. the amino acid isotype of the source tRNA, also showed considerable variability between tissues and between mtDRs and ntDRs [Figure 1G-H]. Nonetheless, certain isotype tRNAs represented a considerable source of tDRs such as Glu, Arg, Gly, and His in ntDRs and Glu, Pro, and Thr in mtDRs, with variations between tissues in terms of the relative abundance of tDRs from these sources.

Analysis of size distribution of detected tDRs in each tissue revealed considerable heterogeneity in ntDRs lengths compared to mtDRs [Figure 2A-B]. In addition, partial least square regression discriminant analysis (PLS-DA) based clustering revealed different clustering patterns between tissue when using ntDRs or mtDRs [Figure 2C-D]. Examining the top 50 variable tDRs, either ntDRs [Figure 2E] or mtDRs [Figure 2F] showed that the spleen and lung tended to cluster together in a distinct cluster compared to other tissues.

**Figure 2:**
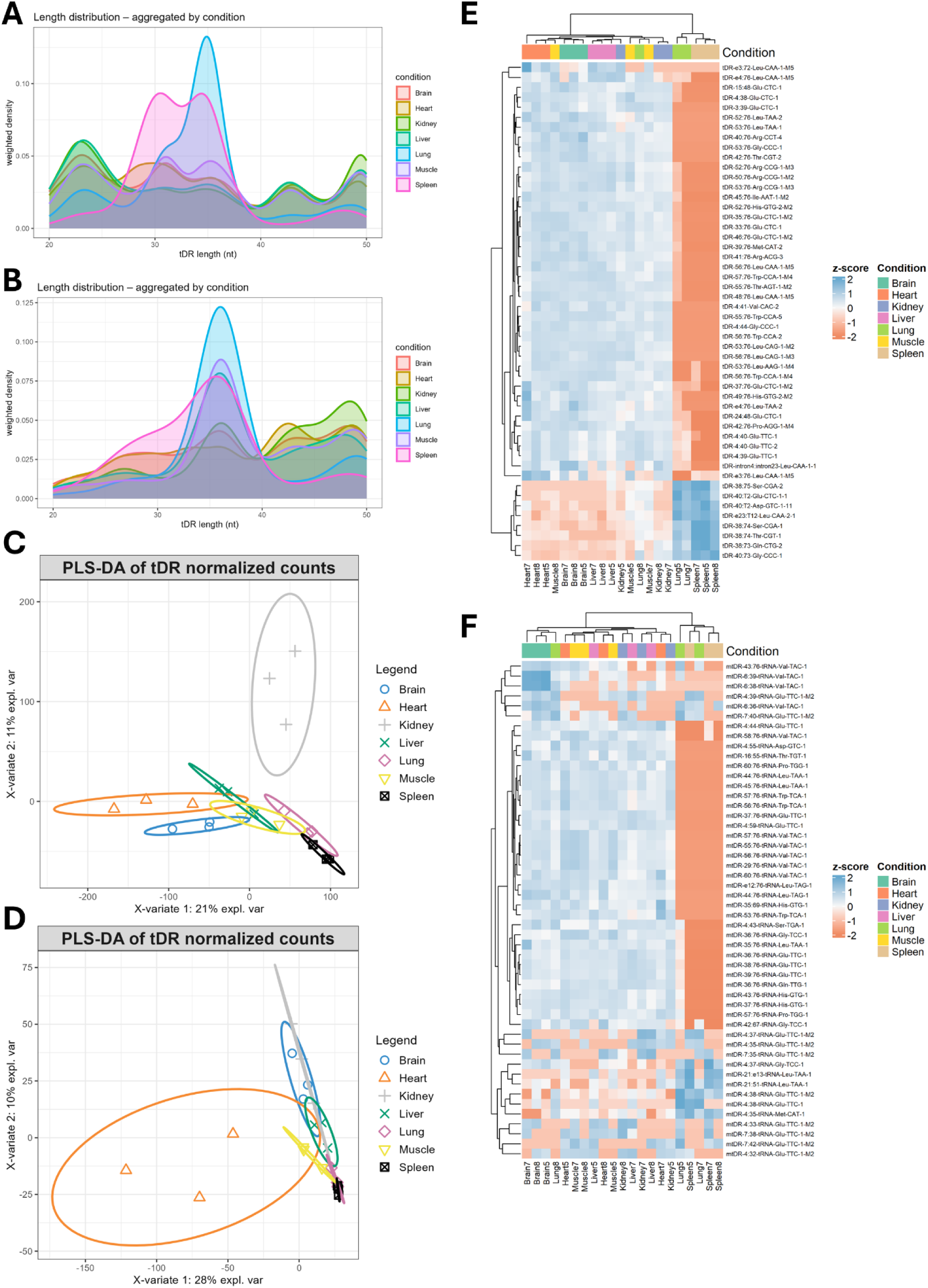
**A:** Length distribution of ntDRs across tissues. **B:** Length distribution of mtDRs across tissues. **C:** Partial least square discriminant analysis (PLS-DA) clustering of ntDRs. **D:** PLS-DA clustering of mtDRs. **E:** Heatmap of top variable ntDRs. **F:** Heatmap of top variable mtDRs.

In summary, tDRs expression reveals inter-tissue heterogeneity in variable aspects of tDRs such as source tRNAs and biogenesis. tDRs expression could also discriminate between different tissues as shown in the clustering data presented.

### tDRs show tissue-specific enrichment patterns

To understand the different patterns of tDRs enrichment in tissues, we conducted differential tDR expression analysis, comparing each tissue to all other tissues (for example, brain vs all, heart vs all, etc.). To avoid the dilution of mtDRs patterns, we analyzed ntDRs and mtDRs separately. Analysis of differentially expressed ntDRs revealed that the tissue most divergent from others was the spleen [Figure 3A-D, supplementary figure 1A-C]. The liver and muscle didn’t show significantly enriched tDRs when compared to all other tissues. Spearman correlation analysis between different tissues using log2FC values of tDRs revealed that the lung, spleen, and kidneys had moderate correlation, with the highest positive correlation between the lung and spleen (Rho = 0.6) [Figure 3E]. The same case was apparent in mtDRs, where spleen showed the highest number of significant tDRs [Figure 3F-H, supplementary figure 1E-I]. However, most tissues did not exhibit significant diversity in their mtDRs expression. Correlation analysis between different tissues using log2FC of mtDRs showed the same correlation between the spleen and lung [Figure 3I].

**Figure 3:**
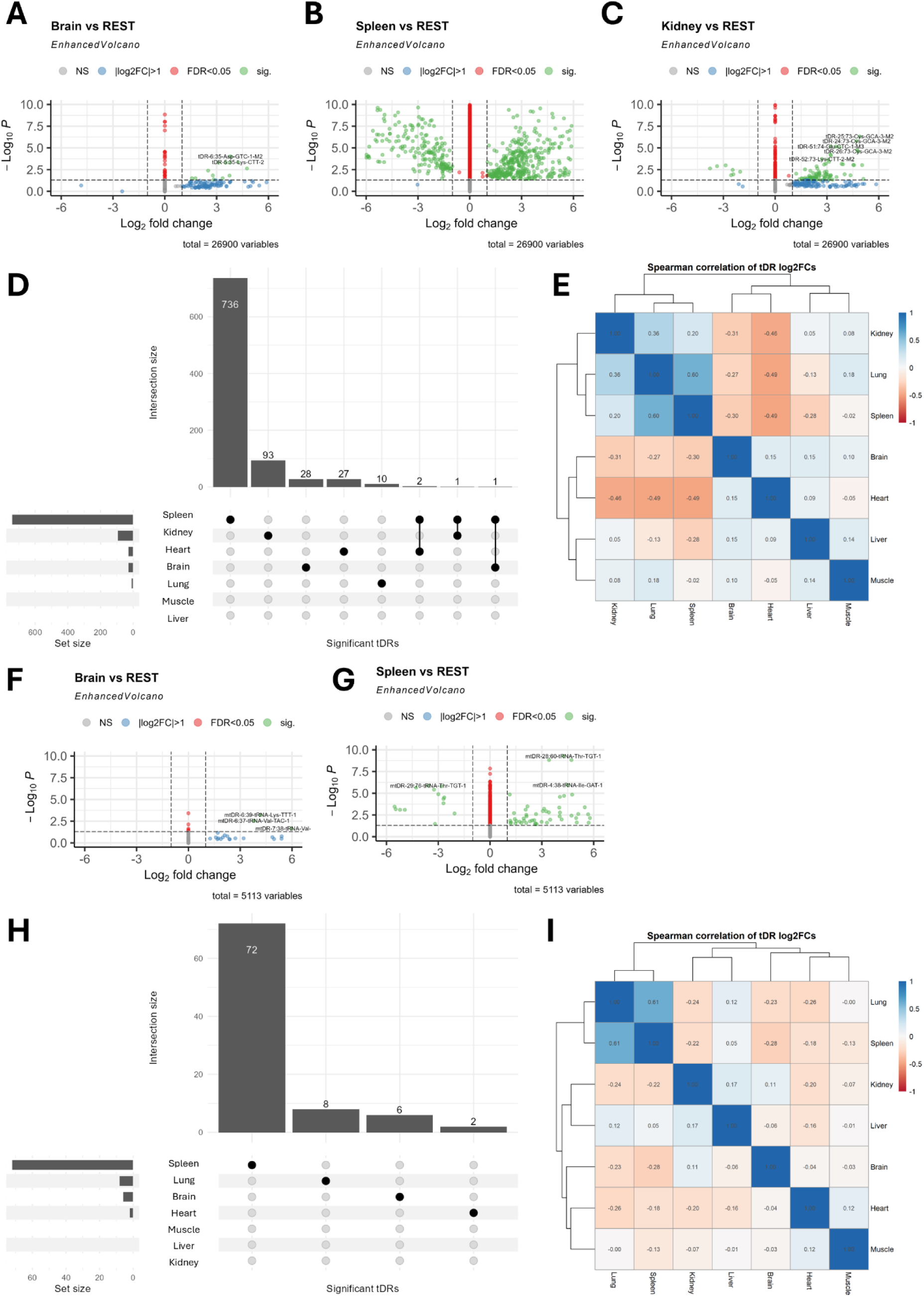
**A-C:** Volcano plot of differentially expressed ntDRs in different tissues. Each volcano represents a tissue vs all other tissues comparison. **D:** Upset plot of differentially expressed ntDRs showing the uniquely expressed ntDRs in different tissues. The spleen had the highest number of uniquely expressed ntDRs. **E:** Spearman’s correlation analysis of differentially expressed ntDRs using their log2 fold change (log2FC) values. **F-G:** Volcano plots of differentially expressed mtDRs in different tissues. **H:** Upset plot of differentially expressed mtDRs showing the spleen with the highest number of uniquely expressed mtDRs. **I:** Spearman’s correlation analysis of differentially expressed mtDRs using their log2FC values.

Next, we explored what tDR subtypes are enriched in the tissue comparisons. Starting with ntDRs, the most abundant differentially expressed class in the brain vs all comparison was 5’tiRNAs followed by i-tRFs, while in the spleen it was tRF3s followed by i-tRFs [Figure 4A-B]. In other comparisons either i-tRFs or tRF3s were the most significantly enriched tDR subclasses [Supplementary figure 2A-C]. The size of upregulated tDRs in significant comparisons showed variations consistent with differences in the relative numbers of enriched tDR subclasses [Figure 4C]. These differences were statistically significant when applying Kolmogorov-Smirnov (K.S.) statistical test [Figure 4D]. Mapping the significantly upregulated tDRs in the tissue vs all comparison to the nucleotide positions in tRNAs shows the heterogeneity in ntDRs origin and differences in biogenesis of tissue specific ntDRs [Figure 4E]. In mtDRs, given that most comparisons yielded few or no enriched tDRs, except for the spleen, we examined the spleen vs all for the tDR subclasses enriched and it was dominated by i-tRFs [Figure 4F]. The brain and lung also showed enrichment of i-tRFs while the heart showed only two enriched tDRs which were of tRF3 subtype [Data not shown].

**Figure 4:**
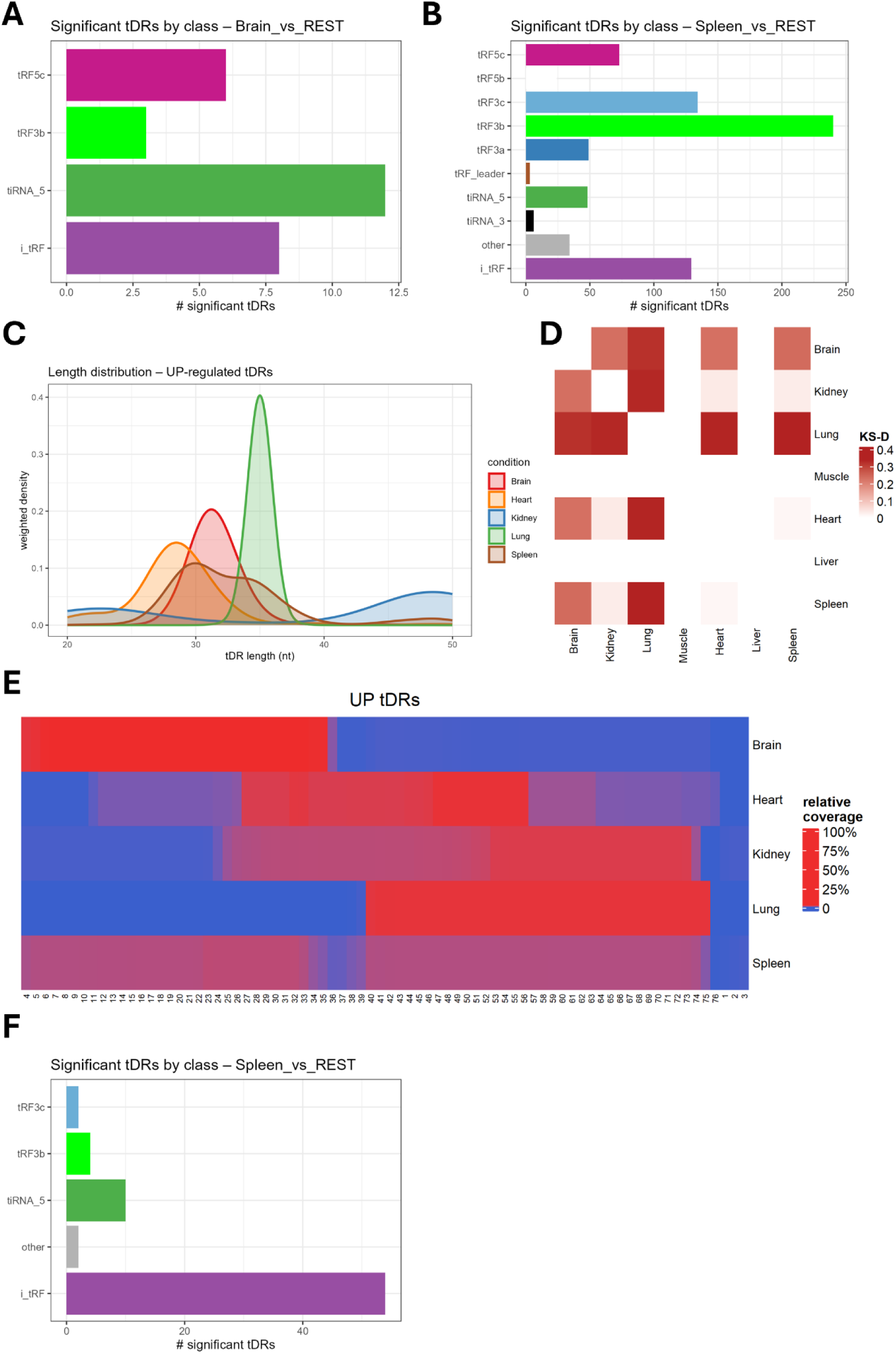
Analysis of the length distribution and subclasses of tissue specific tDRs. **A:** Distribution of tDR subclasses in the upregulated ntDRs in the brain vs rest analysis. **B:** Distribution of tDR subclasses in the upregulated ntDRs in the spleen vs rest analysis. **C:** Size distribution of the upregulated ntDRs in each condition vs rest comparison. **D:** Kolmogorov-Smirnov statistical analysis of length distribution of upregulated ntDRs in each condition. Higher values indicate statistically significant. All comparisons were statistically significant (*p* < 0.05, FDR < 0.05). **E:** Sprinzl heatmap showing the mapping of significantly upregulated ntDRs across the mature tRNA sequences. The heatmap reveals the heterogeneity in tDR subclasses as well as the size distribution differences between tissues.

### tDRs enriched in different tissues have unique tRNA origins

To identify whether ntDRs enriched in different tissues originate from different tRNAs, we identified the isotype (amino acid decoding or isoacceptors tRNA) and the anticodon sequences of the parent tRNAs and conducted Fisher’s exact test with Benjamin-Hochberg false discovery rate (BH-FDR) multiple test correction to identify statistically significant odds that a tDR originate from a specific tRNA isoacceptors [Figure 5A-B]. We observed in the brain that enriched ntDRs were mostly derived from Lys-CTT and Ala-TGC. In the spleen on the other hand, we observed enrichment of ntDRs derived from SeC, Gly, or Val. Other tissues also had different origins of enriched ntDRs [Figure 5A-B]. We next investigated whether specific isodecoder tRNAs could contribute more to the enrichment of ntDRs [Figure 5C, supplementary figure 3A-E]. Indeed, we observed that for each tRNA, several isodecoders had higher odds of being the parent of enriched ntDRs in each tissue. However, some isodecoders were more likely to be the origin of ntDRs than others and importantly, not all isodecoders of a given tRNA were found to be significant in our analysis. Previously, it was reported that tRNA isoacceptors are expressed uniformly in different tissues^48,52^, however, isodecoder expression varied^52^. To test whether mature tRNA isodecoder expression would vary between tissues to explain the enrichment patterns, we re-analyzed the mature tRNA dataset from the same tissues^48^ using the tissue vs all argument [Supplementary figure 4]. In all cases we observed minimal or no statistically significant isodecoder enrichments. Thus, differential enrichment of tRNA isodecoders or isoacceptors cannot explain the differential enrichment of ntDRs in different tissues.

**Figure 5:**
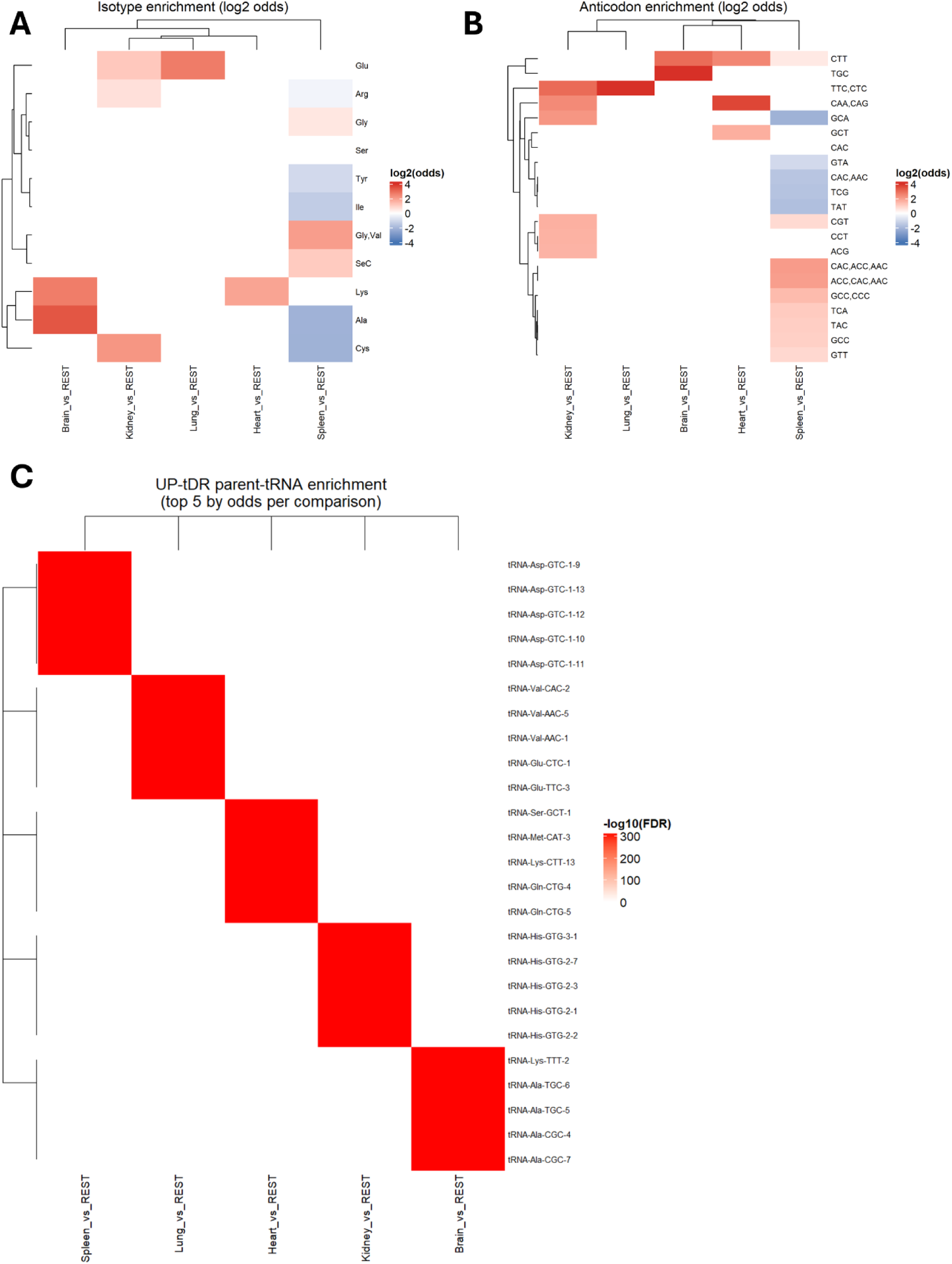
Tissue enriched tDRs have different mature tRNA origins. **A:** Heatmap of log2 Odds ratio of isotype enrichment of upregulated ntDRs. This heatmap indicates whether an isotype tRNA has higher (or lower) odds of producing tDRs in each tissue. **B:** Heatmap of log2 Odds ratio of anticodon enrichment of upregulated ntDRs. This heatmap gives information regarding a given tRNA anticodon (i.e. isoacceptors) being the source of ntDRs in each tissue. **C:** Heatmap of -log10 FDR or parent tRNA enrichment of ntDRs. This heatmap gives information regarding the mature tRNA source of upregulated ntDRs. Importantly, this analysis gives information about whether specific isodecoders are the source of tDRs. The heatmap represents the top 5 tRNAs in each tissue by Odds ratio. Expanded plots are found in supplementary figure 3.

### Highly expressed tissue resident tDRs are heterogenous

As our analysis to this point focused on the tissue specific tDRs, given that we analyzed the upregulated tDRs in the tissue vs all comparisons, we asked whether the highly expressed tDRs by copy number the uniform across tissues. We extracted the top 200 ntDRs by normalized read counts and analyzed them across tissues. We observed variations in tDRs subclasses [Figure 6A] and isotype source [Figure 6B]. For example, tRF5c showed higher enrichment in the spleen while i-tRFs represented the lowest number of tDRs in the spleen but nearly 50% of the top tDRs in the heart and kidney. Heatmap [Figure 6C] of the top 10 expressed tDRs in each tissue and upset plot [Figure 6D] of the top 200 expressed tDRs shows the heterogeneous expression of highly expressed tissue resident tDRs across tissues. Importantly, the spleen and lungs closely clustered together [Figure 6C], following the trend observed at multiple analysis levels that was also previously reported at the level of mRNA translation and tRNA modifications^48^. Furthermore, analysis of the isotype tRNA and isoacceptors tRNA sources of the top 200 tDRs yielded varying enrichment patterns across tissues [Supplementary figure 5]. For example, the spleen and lungs showed enrichment of tDRs derived from Asp-GTC. On the contrary, all tissues showed significant enrichment of tDRs from Gln tRNAs except for the lung and spleen. Overall, our analysis reveals significant heterogeneity in terms of tDRs expression across different tissues as well as the source of those expressed tDRs and their biogenesis.

**Figure 6:**
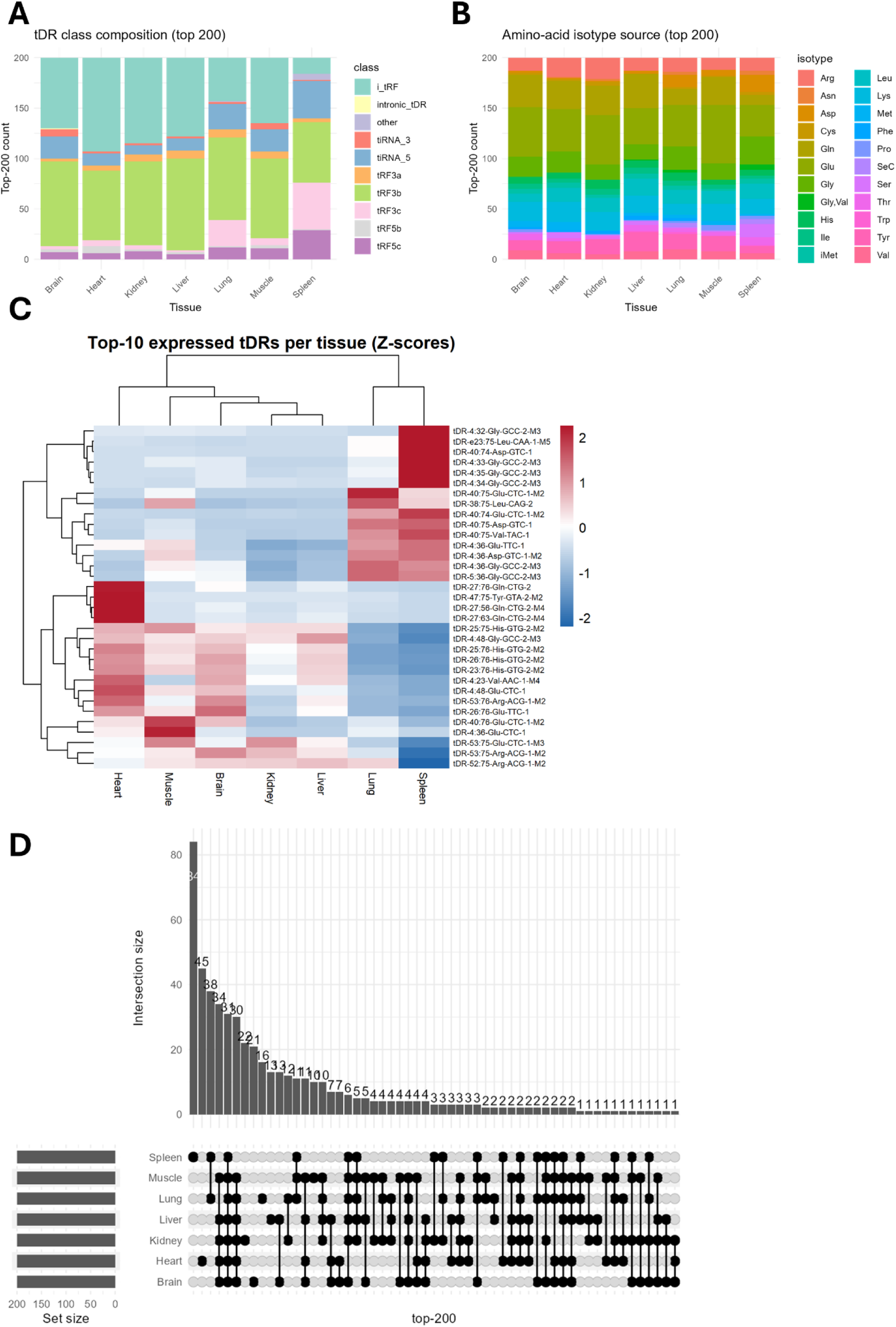
The highest expressed tDRs are heterogenous across tissues. **A:** Subclasses of highly expressed (top 200 by normalized read counts) ntDRs in different tissues. **B:** isotype classes of highly expressed ntDRs in different tissues. **C:** Heatmap of the top 10 expressed ntDRs in each tissue. **D:** Upset plot of the top 200 expressed ntDRs showing overlapped and unique ntDRs across tissues.

### G4 quadruplexes forming tDRs are enriched in the spleen

tDRs functions have come under significant scrutiny in recent years^53^. tDRs were proposed to interact with RBPs via motif sequences to displace them from their binding mRNAs^26^, to enhance translation of ribosomal mRNAs^28,29^, and to form G4 quadruplexes^17^. Given that in RNA function follows form^1,35^, we interrogated whether tissue resident and enriched ntDRs can form G4 quadruplexes. We used pqsfinder^54^ to predict whether a given tDR is able to form G4 quadruplex or not. First, we analyzed the top 200 enriched ntDRs in each tissue. We observed varying numbers of potential G4 quadruplexes forming ntDRs in different tissues, with the highest numbers occurring in the spleen and lungs [Figure 7A]. However, the pqsfinder score, which predicts the strength of the potentially formed G4 structure^54^, was more or less evenly distributed across tissues. We examined the isotype origin of the predicted G4 quadruplexes forming tDRs and their tDR subclass and found that most of these tDRs in all tissues originate from Glycine tRNAs [Figure 7C]. The lungs and spleen showed a population of Asp, Val, and SeC originating tDRs also capable of forming G4 quadruplexes that did not show or only had minor contribution in other tissues. Other tRNAs such as Gly, Leu, and Arg also contributed with varying extents in different tissues. Examining the tDR subclasses that contribute to G4 quadruplex formation revealed more heterogeneity between tissues, especially when considering spleen and lungs [Figure 7D]. While most tissues showed that the G4 quadruplexes forming ntDRs to be of the i-tRF subtype, the spleen and lungs showed more contribution for tRF3 and to a lesser extent (but higher than other tissues) 5’tiRNAs.

**Figure 7:**
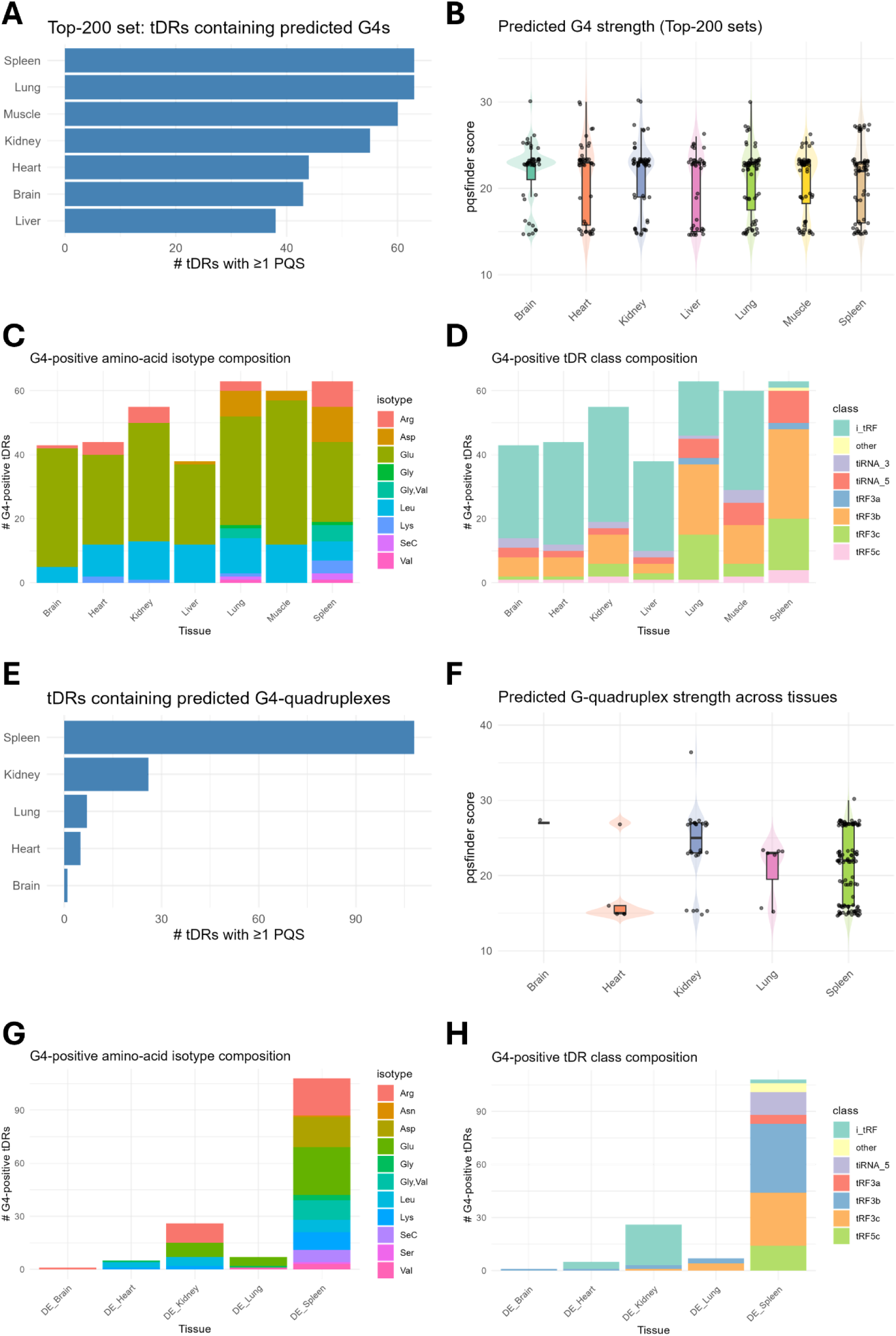
ntDRs can form G4 quadruplexes with special enrichment of G4 forming tDRs in the spleen. **A:** Bar plot showing the numbers of ntDRs predicted to be able to assemble into G4 quadruplexes in each tissue from the top 200 expressed ntDRs (top ntDRs). **B:** pqsfinder score of the potentially G4 forming top ntDRs in each tissue. **C:** Isotypes of the G4 forming top ntDRs in each tissue. **D:** tDR subclasses of the G4 forming top ntDRs. **E:** Bar plot representing the numbers of predicted G4 forming ntDRs in the tissue enrichment analysis (i.e. differentially expressed ntDRs). **F:** pqsfinder score of the potentially G4 forming differentially expressed ntDRs in each tissue. **G:** Isotypes of the G4 forming differentially expressed ntDRs in each tissue. **H:** tDR subclasses of the G4 forming differentially expressed ntDRs.

Analysis of the differentially expressed ntDRs (i.e., those with specific tissue enrichment) revealed that the spleen is specifically enriched in potential G4 forming ntDRs followed by the kidneys [Figure 7E-F]. Examining the isotype and class distribution of G4 quadruplexes forming differentially expressed ntDRs revealed a different picture compared to the top expressed ntDRs [Figure 7G-H]. For example, there was a more uniform distribution of the isotype contribution to the G4 forming ntDRs in the spleen with less dominance of the Glu originating tDRs [Figure 7G]. tRF3 was the most dominant subclass in the G4 forming ntDRs in the spleen while in other tissues, such as kidneys, i-tRFs were the dominant class [Figure 7H].

In summary, tDRs highly expressed and enriched in different tissues show a potential to form G4 quadruplexes. While previously only stress-induced 5’tiRNAs were the subject of research focusing on G4 quadruplexes-forming potential of tDRs^17,31^, other subclasses of ntDRs show a potential for G4 quadruplexes assembly. Nonetheless, this remains a computational prediction and more biochemical and molecular analysis are needed for validation.

## Discussion

In this analysis, we reveal the significant heterogeneity in nuclear and mitochondrial tDRs across different tissues. Importantly, tDRs are heterogeneous at many levels including source tRNAs, size, site of cleavage/biogenesis, subclass, structure, and potential tRNA modifications in their sequences from the source tRNAs. We also noted that while the spleen had the lowest number of detected tDRs compared to other tissues, it was the most diverse. tDRs were shown to be important regulators of post-stroke immune response and can replace microRNAs to regulate immune response^55^. The closest tissue to spleen was the lungs, which are also known for their high macrophage and immune cell populations. The brain, which is translationally and epitranscriptionally unique compared to other tissues^48^ showed preference for the enrichment of 5’tiRNAs, compared to the higher i-tRF or tRF3 enrichments in other tissues. Source tRNAs also showed significant heterogeneity across tissues. Collectively, this information indicates that the biogenesis and production of tDRs in different tissues is not a matter of simple tRNA turnover, but a highly orchestrated process that potentially serves to regulate gene expression and physiological functions of different tissues and cells. A potential example for this specialized functionality of various tDRs is the enrichment of potentially G4 quadruplexes forming ntDRs in the spleen. G4 quadruplexes have been shown to regulate tDRs function via facilitating phase separation or RBPs binding^17,31,56^. The enrichment of potentially G4 quadruplexes forming ntDRs in the spleen alludes to a potential immune function of this subset of tDRs that was not previously studied.

It is important to note that to fully understand tDRs biogenesis and function, given their extreme heterogeneity, more datasets, and importantly more robust datasets, are needed. In addition to robust tDR profiling, combining tDR sequencing with approaches to map tRNA or tDR modifications, G4-quadruplexes formation, or potential RBPs bindings at a massive scale, will be of essence in deciphering the functions of tDRs as well as the rules of their production. To date, the biogenesis of tDRs is not fully understood. While several enzymes apart from Angiogenin have been shown to play important roles in cleaving tRNAs and in turn regulating mRNA translation via codon biased translational regulation^9,11,57^, more work is needed to correlate tDRs with specific translational phenomena. On the global scale, the tDRs expressed in each tissue do not correlate directly with specific codon decoding patterns. While not shown here, comparing each tissue to all others using Ribo-seq data from the same tissues and study^48^, A-site, P-site, or E-site ribosome dwelling times did not match the anticodons of the tDRs source tRNAs. Thus, it appears that the regulation of codon biased translation by directed tRNA cleavage is a context/condition specific phenomenon^9,11,57^ that occurs under stress conditions. Again, such discrepancies highlight the shortcomings in our understanding of tDR biology.

In conclusion, tDR biogenesis is extremely heterogeneous under physiological conditions but appears to follow specific rules. Mature tRNA expression, either at the isoacceptors or isodecoders levels, does not appear to be a contributing factor to which tRNAs produce the most tDRs. What are the rules regulating tDR production? Do G4 quadruplex forming tDRs play specific roles in the immune response or immune cells? Such questions, and many more, need to be addressed in future work to fully comprehend tDR biology, understand their roles in the many diseases they were reported to play a role in, and to potentially aid in developing novel tDR-based therapeutics.

## Methods

**Datasets:** small RNA-seq dataset was retrieved from a previously published study^48^ from sequence read archive project number PRJNA1003133

### Data analysis

tDRnamer standalone version^8^ (https://trna.ucsc.edu/tDRnamer/docs/standalone/) was used to analyze tDRs in the small RNA-seq dataset after adaptor trimming and collapsing paired end reads to single fastq.gz file using SeqPrep (https://github.com/jstjohn/SeqPrep) using max sensitivity, fastq.gz files, and mm39 database (or mitochondrial mm39 for mtDRs). tDRs with fewer than 10 reads were excluded. We used the output tDR-info.txt file for our downstream analysis.

To analyze tDRnamer output, we used a custom R script that first gathered the read counts for detected tDRs to create a count matrix as well as an annotation file containing all the tDRs information such as start and end positions, source tRNAs, etc. tDRs were filtered to include those between 15 and 50 nucleotides. DESeq2^58^ was used for differential tDR enrichment analysis as well as generation of normalized tDR count files for downstream analysis. Upset plots were generated using ComplexUpset package in R. PLS-DA analysis was conducted using mixOmics^59^ package in R. Classifying tDRs into subclasses was conducted using a custom R script based on the currently accepted tDR classifications. Analysis of length distribution was done using the information provided by tDRnamer. Weighted Kolmogorov–Smirnov test to compare between different conditions for statistical significance in enriched tDRs lengths. Sprinzl heatmap was generated using the start and end positions of each tDR provided by tDRnamer. To identify the source of tDRs, isotype, anticodon, and tRNA copy (isodecoder) information derived from tDRnamer output was used for Fischer’s exact test with Benjamin-Hochberg false discovery rate (FDR) multiple testing correction. Odd ratio and FDR were used to identify significance and generate plots. G4 quadruplexes analysis was conducted using pqsfinder^54^. All heatmaps, volcano plots, and other visualizations were created in R.

## Acknowledgments

The authors report no financial or ethical conflicts of interest regarding this work.

## Author contributions

**D.A**: Writing/Editing. Data review. **S.R**: Conception and study design. formal Data analysis. Scripts design. Funding. Supervision. **K.N**.: Critically reviewed the manuscript. All authors approved the final version of the manuscript.

## Funding

This work was supported by The Japan society for the promotion of science (JSPS) kakenhi grant number 23H02741

## Supplementary figure

**Supplementary figure 1 (supplement to Figure 3):**
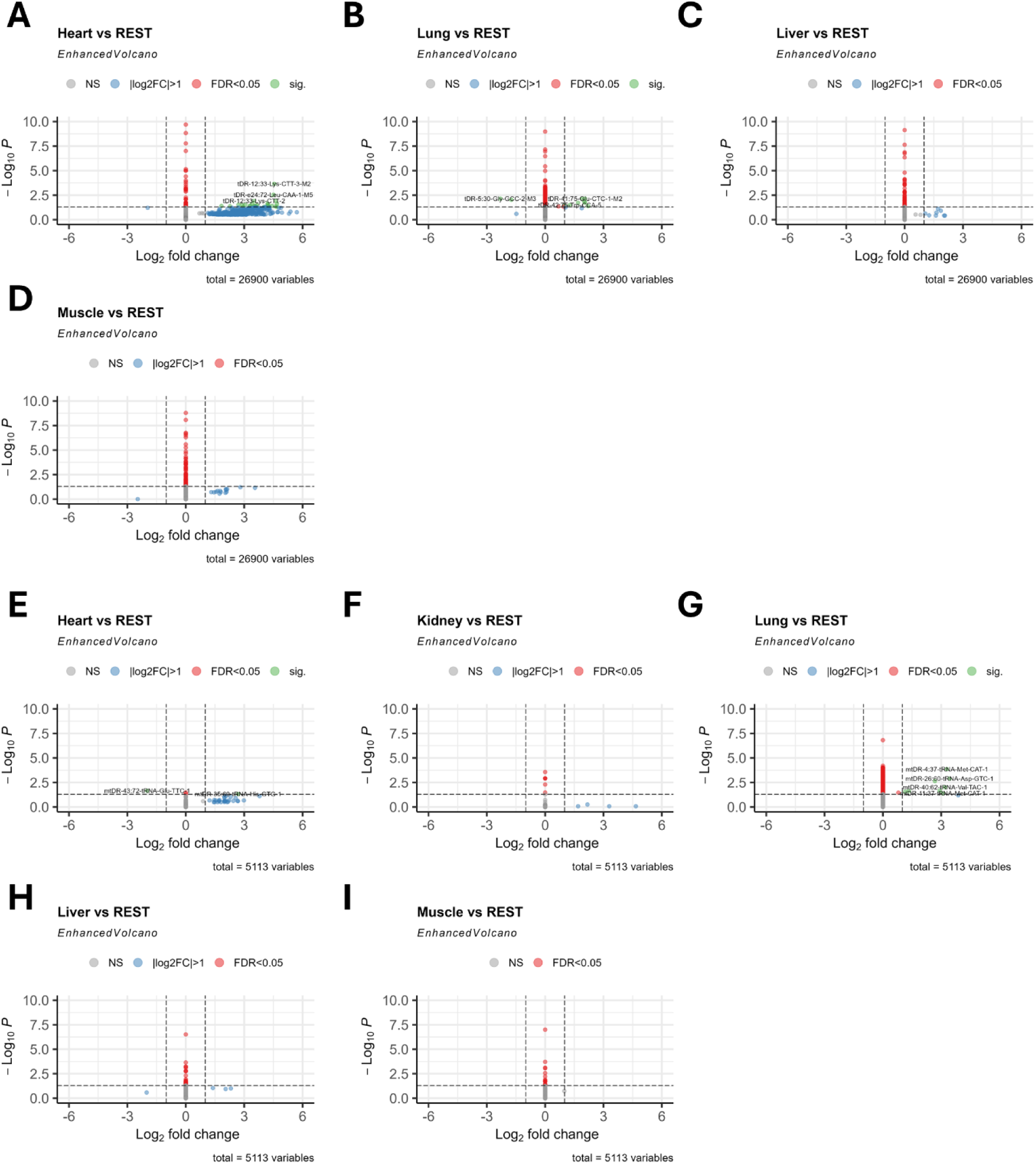
**A-D:** Volcano plot of differentially expressed ntDRs. **E-I:** Volcano plot of differentially expressed mtDRs.

**Supplementary figure 2 (supplement to figure 4).**
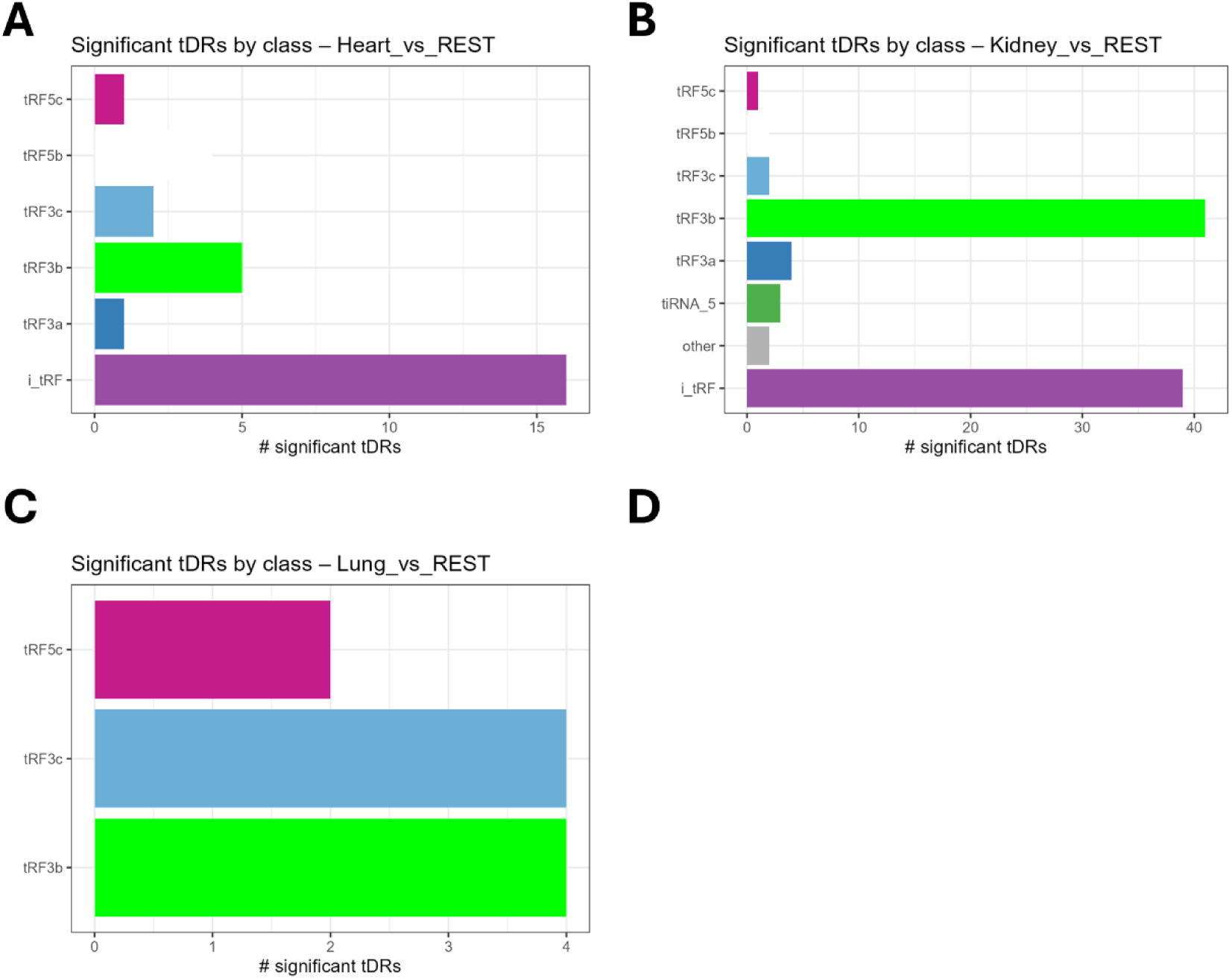
**A-C:** Distribution of tDR subclasses in the upregulated in different tissues.

**Supplementary figure 3 (supplement to figure 5):**
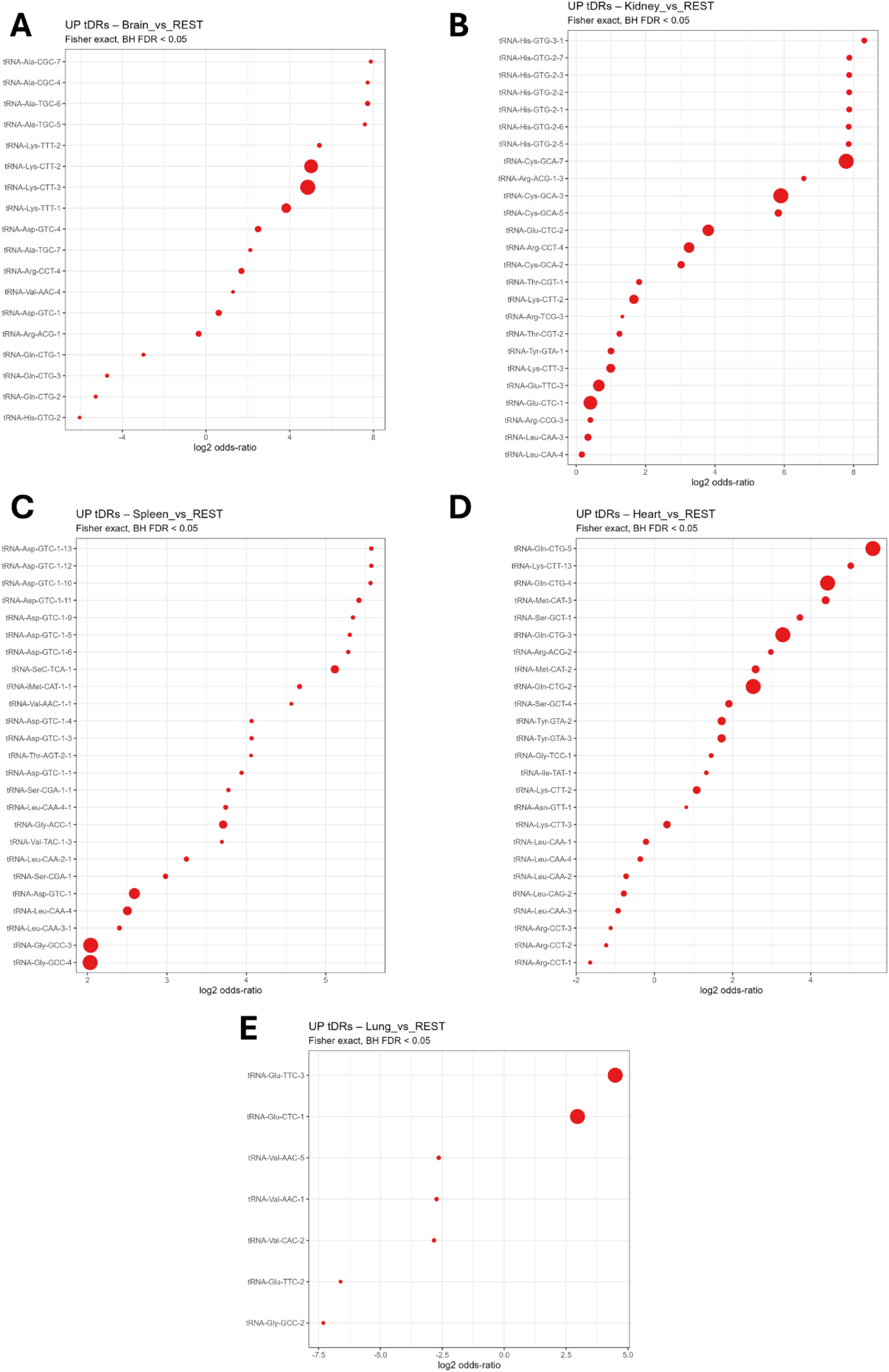
Dot plot of log2 odds ratio of an isodecoder tRNA to be the origin of ntDRs in different tissues. The size of the bubble reflects the number of enriched ntDRs.

**Supplementary figure 4:**
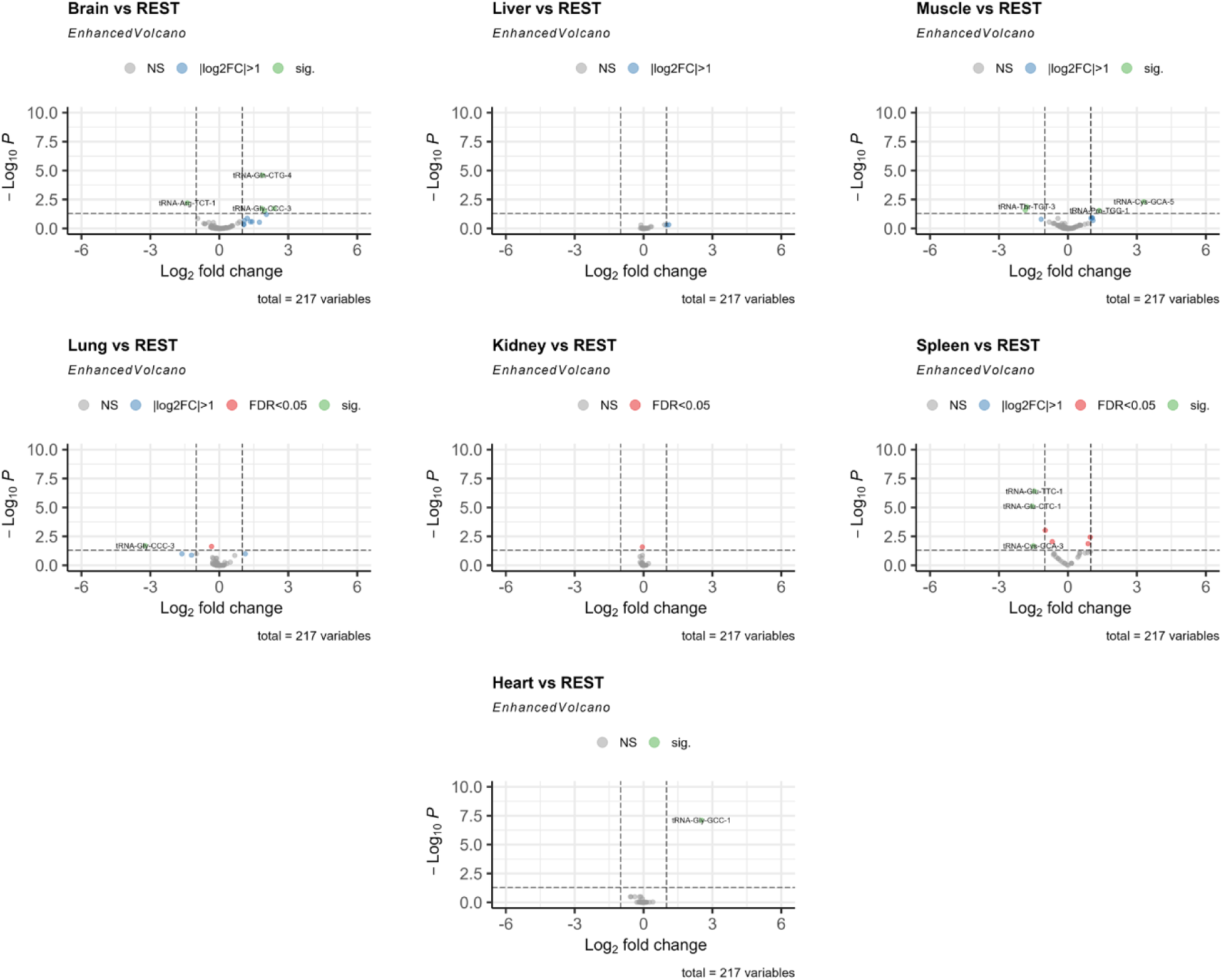
Analysis of mature tRNA expression in different tissues. Here, we conduct differential mature tRNA expression analysis comparing each tissue to all other tissues (as done for tDRs). The volcano plots presented herein reveal minimal to no changes in tRNA expression across tissues.

**Supplementary figure 5:**
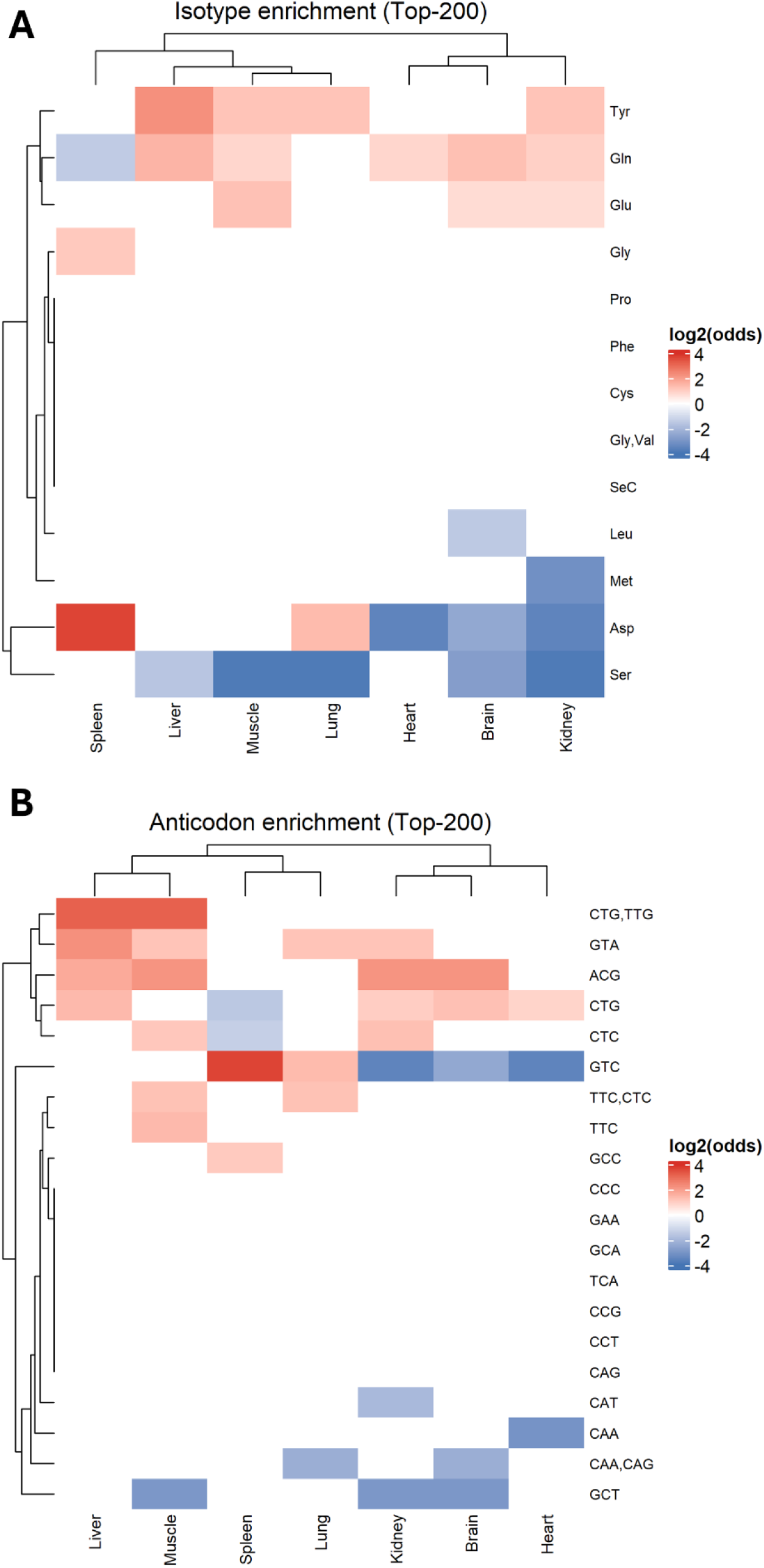
Isotype enrichment (**A**) and Anticodon (**B**) enrichment of the top 200 ntDRs in each tissue showing variations in source tRNAs with unique signatures in the spleen and lungs. The heatmaps represent log2 Odds ratio derived from Fisher’s exact test with Benjamin-Hochberg (BH) multiple test correction analysis.

## Notes

### Competing Interest Statement

The authors have declared no competing interest.

